# The Shexiang Baoxin Pill Protects Myocardial Cells from Multiple Targets of MIRI through the PI3K/Akt/eNOS Signal Pathway

**DOI:** 10.1101/2023.11.13.566957

**Authors:** WEI Na, LI Siyuan, GAO Yuan, LIU Zhenbing

## Abstract

**Background:** Myocardial ischemia-reperfusion injury (MIRI) can significantly aggravate myocardial injury in patients with ST-segment elevation myocardial infarction (STEMI). At present, there are few effective treatments for MIRI. The Shexiang Baoxin Pill (SBP) can reduce MIRI. The PI3K/Akt/eNOS signaling pathway, inflammation, oxidative stress, and apoptosis are all involved in the regulation of MIRI. SBP has multi-component, multi-target, and synergistic effects, but its mechanism of action on MIRI has not been reported.

**Purpose:** We sought to explore whether SBP exerts a protective mechanism by inhibiting the inflammatory reaction, oxidative stress, and apoptosis, reducing MIRI through the PI3K/Akt/eNOS signal pathway.

**Materials and methods:** Hypoxia-reoxygenation (H/R) H9c2 cardiomyocytes were used as an in vitro model of MIRI. The active components of Shexiang Baoxin pills were extracted with water. The levels of phosphorylated proteins and genes related to the PI3K/Akt/eNOS pathway were measured by Western blotting and real-time fluorescence quantitative PCR. Cell viability, apoptosis rates, and apoptosis-related proteins (Bcl-2, Bax, Caspase-3) were detected by CCK-8, flow cytometry, and Western blotting. The expression of reactive oxygen species (ROS), homocysteine (Hcy), malondialdehyde (MDA), and gp91^phox^ was detected by fluorescence probe, ELISA, TBA, and Western blotting. The levels of inflammatory factors (TNF-α, IL-6, IL-18) were measured by an ELISA method.

**Results:** SBP increased the cell survival rate of H/R cardiomyocytes, reduced the injury to H/R cardiomyocytes, and increased the protein phosphorylation levels of p-PI3KY607, p-AktSer473, p-eNOSSer1177, and mRNA of H/R cardiomyocytes. In addition, SBP increased the level of Bcl-2 protein and the Bcl-2/Bax ratio and decreased the apoptosis rate and Bax and Caspase-3 expression. It reduced the levels of oxidative stress indexes (ROS, HCY, MDA, and gp91phox) and inflammatory factors (TNF-α, IL-6, IL-18) and enhanced antioxidant stress, anti-apoptosis, and an anti-inflammatory reaction. The above effects were attenuated after the inhibition of the PI3K/Akt/eNOS signal pathway.

**Conclusion:** We established that SBP extract inhibited oxidative stress, inflammatory response, and apoptosis through the PI3K/Akt/eNOS signal pathway and alleviated the injury of H9c2 cells induced by hypoxia-reoxygenation.

## 1. Introduction

Acute myocardial infarction (AMI), one of the most serious cardiovascular events, is still the main cause of morbidity and mortality in the world [1]. ST-segment elevation myocardial infarction (STEMI) is the most severe type of acute myocardial infarction. In Europe, more than 500,000 STEMI patients are hospitalized each year. Timely thrombolysis and percutaneous coronary intervention are the most effective methods to reduce the extent of myocardial infarction and improve clinical prognosis [2]. However, myocardial ischemia-reperfusion injury (MIRI) can be induced after myocardial reperfusion, which in turn leads to cardiomyocyte death and an increase in the myocardial infarction area [3]. MIRI is one of the main factors affecting the clinical outcomes in STEMI patients [4], and in severe cases, it can lead to death [5–6]. Although some treatments, such as ischemic adaptation, drug therapy, and physical methods (hyperoxia, hypothermia, and nerve electrical stimulation), can reduce MIRI [7–8], the efficacy of these single therapies and single-targeted drug therapies on MIRI has not been established [9–12]. The pathogenesis of MIRI is highly complex, involving multiple targets. Therefore, a new method of multi-target therapy is urgently needed to reduce the risk of MIRI [13].

The pathogenesis of MIRI is complex and diverse and includes oxidative stress, ischemia and hypoxia, calcium overload, and inflammatory reactions[14]. In 2019, the European Union for Heart Protection put forward the principle of multi-target therapy for MIRI. For example, intracellular targets can include therapeutic strategies such as survival-promoting signaling pathways, cell death pathways, and organelles. The reasonable combination of different protection strategies will help to activate complementary survival pathways and/or inhibit harmful cell death pathways, resulting in improved cardioprotective effects [15]. Animal experiments have recently confirmed that ischemic preconditioning protects against MIRI in diabetic rats by activating the serine/threonine protein kinase (Akt)-related signaling pathway and inhibiting apoptosis[13]. Therefore, multi-target therapy for MIRI is a novel approach for its treatment[15]. In particular, oxidative stress, inflammation, and apoptosis play critical roles in the onset and development of MIRI [16]. It has been confirmed that the phosphatidylinositol 3 kinase (PI3K)/Akt/ endothelial nitric oxide synthase (eNOS)-signaling pathway is involved in the regulation of oxidative stress, inflammation, and apoptosis in MIRI [17–18].

The Shexiang Baoxin Pill (SBP) is one of the most commonly used Chinese medicines in the treatment of AMI over the past 40 years in China [19–20]. Qin et al. found that SBP reduced MIRI and myocardial infarction size in patients with STEMI [21]. Several studies have also shown that SBP can reduce MIRI in experimental rats[21–22], as the extracted components of SBP can reduce MIRI and infarct size by inhibiting oxidative stress, inflammatory reactions, and apoptosis[23–24]. However, there is no existing report that describes the mechanism of the multi-target action of SBP on MIRI. Therefore, in this study, we developed an H9c2 hypoxia-reoxygenation (H/R) cardiomyocyte model to explore whether SBP exerts anti-apoptotic, anti-oxidative stress, and anti-inflammatory effects and provides multi-target protection against MIRI through the PI3K/Akt/eNOS signal transduction pathway.

## 2. Materials and methods

### 2.1 Cell culture

We purchased the H9c2 (2-1) rat cardiomyocyte line from the Cell Resource Center of Shanghai Academy of Life Sciences, Chinese Academy of Sciences,Shanghai,China. Cell culture reagents were from Seamer Fisher Technologies, Waltham, MA, USA. Cardiomyocytes were cultured in Dulbecco’s modified Eagle medium (DMEM) supplemented with 10% fetal bovine serum, 100 µg/mL streptomycin, and 100 U/mL penicillin. The cells were cultured in a humidified environment containing 5% carbon dioxide in compressed air at 37°C. When the degree of cell fusion was over 80%, the cell clumps were digested with trypsin and passaged, and the cells with favorable development and in a logarithmic growth phase were selected for follow-up experiments.

### 2.2 Reagents

SBP was purchased from Shanghai Hehuang Pharmaceutical Co., Ltd., with a specification dose of 22.5 mg × 42 pills/bottle (Chinese medicine standard code, Z31020068; batch number, 210405). The SBP was ground into a powder and dissolved in distilled water to obtain a suspension of 50 mg/mL, which was diluted to 2 mg/mL SBP in DMEM for further use. CCK-8 kit, trypsin, BCA kit, ECL luminous solution, embossing cassette, X-OMAT BT film, electrophoresis solution, PVDF film, lysate, and TBST were purchased from Shanghai Biyuntian Biotechnology Co., Ltd. High-sugar DMEM medium, sugar-free DMEM medium, and fetal bovine serum (FBS) were purchased from Gibco Company, USA; penicillin-streptomycin from Hyclone Company, USA; methanol and dichloromethane from Sinopharmaceutical Group; the Annexin-V FITC/PI double-staining kit from the BD Company; and transmembrane solution, bovine serum albumin (BSA), and rainbow 180 broad-spectrum protein marker from Beijing Solebao Biotechnology Co., Ltd. P-PI3K antibody (1:500, rabbit anti, ab182651), eNOS antibody (1:1000, rabbit anti, ab199956), Bcl-2 antibody (1:1000, rabbit anti, ab194583), and gp91^phox^ (1:2500, rabbit anti, ab129068) were purchased from Abcam company; PI3K antibodies (1:1000, rabbit anti, 4257S), Akt antibodies (1:1000, rabbit anti, 4691s), p-Akt antibodies (1:500, rabbit anti, 4060S), p-eNOS antibodies (1:500, rabbit anti, 9570S), Bax antibodies (1:1000, rabbit anti, 14796S), caspase-3 antibodies (1:1000, rabbit anti, 14220S), β-actin antibodies (1:1000, rabbit anti, 4970S), and goat anti-rabbit secondary antibodies (1:2000, 7074S) were purchased from the CST company.

### 2.3 Establishment of the hypoxia-reoxygenation model and groups

H9c2 cells were inoculated in 96-well plates at a density of 5000/well. After 24 h of culture, high-sugar DMEM was replaced with sugar-free, serum-free DMEM medium, cells were exposed to simulated hypoxia for three hours (at an oxygen concentration during hypoxia of 0%), and the medium was replaced with high-glucose DMEM reoxygenation for two hours (the oxygen concentration during reoxygenation was 95%). The cells were randomly allocated to nine groups: a control group, normal cultured cardiomyocytes with the same volume of high-glucose DMEM; Hmax R group; 2-mg/mL SBP group—cardiomyocytes pretreated with high-glucose DMEM containing corresponding concentrations of SBP for 24 h; H/R+LY294002, H/R+MK2206, and H/R+L-NAME groups pretreated with LY294002 (10 µM), MK2206 (5 µM), or L-NAME (200 µM) for 25 h before treatment, respectively; and H/R+LY294002+SBP, H/R+MK2206+SBP, H/R+L-NAME+SBP groups. Before treatment with Hacher, the rats were pretreated with LY294002 (10 µM), MK2206 (5 µM), or L-NAME (200 µM) for one hour [25–27], and then each group was pretreated with 2 mg/mL [28] SBP for 24 h.

### 2.4 Cell counting kit-8 (CCK-8) assay

The corresponding treated cells of each group were combined with 10 µL CCK-8 reagent, gently shaken, and incubated at 37°C for two hours. The optical density (OD) at a 450-nm wavelength was measured by an enzyme-labeling instrument(BioTek,ELX-800,Vermont,USA), and the cell survival rate (%) = (experimental group OD value – blank group OD value)/(control group OD value – blank group OD value) × 100%.

### 2.5 Flow cytometry

For apoptosis analysis, H9c2 cells were inoculated in six-well plates, and the cells were treated with the corresponding conditions in each group. After the intervention, the cells were digested with trypsin without EDTA, collected, centrifuged at 1200 × g for 5 min at 4°C, washed twice with precooled PBS, and centrifuged again for 5 min at 1200 × g. Cardiomyocytes were resuscitated with a buffer solution, the cell density was adjusted to 1 × 10^6^ cells/mL, and 100 µL of cells was used for staining. We added 5 µL of annexin V-FITC and 10 µL of PI and mixed the solution gently for 10–15 min protected from light and reacted at room temperature. Binding buffer (400 µL) was added, and the solution was mixed and placed on ice and analyzed with a FACSCalibo flow cytometer (BD Bioscience, Franklin Lake, NJ, USA) within one hour, and the results were analyzed with the BD CellQuest pro Software. The apoptotic rate (%) = early apoptotic rate + late apoptotic rate.

### 2.6 Fluorescence-probe methodology

H9c2 cells with good growth were randomly allocated to six-well plates and inoculated with 1.5 × 10^5^ cells per well. After the successful establishment of the model, the DMEM in the six-well plate was aspirated with a pipette, and 1 mL of 10 µmol LDCFH-DA was added and diluted with serum-free culture medium and incubated for 20 min. The cells were then washed three times with serum-free DMEM to remove DCFH-DA that did not enter the cells. Finally, we directly observed the fluorescence intensity and recorded it in real-time by fluorescence microscopy(BD FACSCalibur,USA) at a 488-nm excitation wavelength and 525-nm emission wavelength, and the mean fluorescence intensity was analyzed with Image J (NIH, USA).

### 2.7 Determination of malondialdehyde (MDA) content in cells using the TBA method

Malondialdehyde (MDA) is a degradation product of lipid peroxidation and can be condensed with thiobarbituric acid (TBA) to form a red product with a maximal absorption peak at 532 nm. Because the substrate here is TBA, this method is called the TBA method. The treated cells of each group were scraped with cell curettage and transferred to a 1.5-mL centrifuge tube. The fifth extract from the intracellular malondialdehyde kit (0.5 mL) was added and mixed for two min. The cells were broken by ultrasonic fragmentation, creating a suspension. The suspension samples of each group were sampled at 0.1 mL from the 1.5-mL centrifuge tube, the same volume of anhydrous ethanol was added to the tube, and the same volumes of 10 nmol/mL standard and from the 1.5-mL centrifuge tube were added to the standard tube. Finally, the working liquid from the MDA kit in the 1-mL cell prep was added to all the tubes, and the needle was heated with an alcohol lamp in advance to pierce a small hole in the tube cover; we then covered the tube and vortexed the solution in a water bath for 40 min above 95°C. After removal, the tubes were cooled with running water and centrifuged at 4000 × g for 10 min. The empty plate was scanned at 490 nm. The reaction solution of each tube of 0.25 mL was accurately absorbed onto the 96-well plate that had just been scanned, and the absorbance of each hole was determined. MDA (nmol/mg prot) = (measured OD value – blank OD value)/(standard OD value – blank OD value) × standard concentration (10 nmol/mL)/sample protein concentration (mg prot/mL).

### 2.8 Enzyme-linked immunosorbent assay (ELISA)

After the corresponding treatment, the cells of each group were collected, suspended, and centrifuged at 3000 × g for 15 min. Then, the supernatant was removed, 1 mL of PBS was added, and the mixture was placed at –20°C overnight. After repeated freeze-thaw treatment to destroy the cell membranes, the supernatant was prepared by centrifuging at 5000 × g for 5 min at 4°C. We then set up the standard hole and the sample hole to be tested, added 100 µL of the sample or the standard to each hole, covered the holes with plate paste, and incubated it at 37°C for two hours. The liquid was then discarded, the plate was dried, and 100 μL of biotin-labeled antibody-working solution was added to each hole, covered with plate paste, and incubated at 37°C for 1 h. We discarded the liquid and shook the prep dry, add 200 μ L/hole of washing solution,soaked it for 2 min,and shook the prep dry,washed the plate three times. We added 100 μL of horseradish peroxidase-labeled avidin solution to each hole, covered them with plate paste, and incubated at 37°C for 1 h. We discarded the liquid and shook the prep dry, add 200 μL/hole of washing solution,soaked it for 2 min,and shook the prep dry, washed the plate five times. Ninety microliters of substrate solution was added to each well in order, covered with plate paste, and incubated for 20 min at 37°C. Terminator (50 μL) was then added to each well to terminate the reaction. Finally, the OD value of each well was measured sequentially at an internal 450-nm wavelength at 5 min.

### 2.9 Western-blot analysis

Differentially treated H9c2 cells were collected from the six-well plates and lysed for 40 min in a mixture of RIPA buffer and protease/phosphatase inhibitor (Beyotime, Haimen, China). The lysate was centrifuged at 12000 × g for 20 min, leaving only the supernatant as the total protein. The protein concentration was determined by a BCA protein kit (Beyotime, Haimen, China), and the protein sample was diluted in 4 × SDS sample buffer, heated to 100°C for 5 min, and separated in a 10% SDS-polyacrylamide gel. The protein was transferred to PVDF membranes and incubated in TBST containing 5% bovine serum albumin for two hours at room temperature. We incubated the PVDF film with the first antibody overnight at 4°C, washed extensively in TBST containing 0.1% Tween-20, incubated with the second antibody coupled with HRP at room temperature for one hour, and washed again with TBST containing 0.1% Tween-20. The PVDF film was infiltrated with ECL photoluminescence solution, placed away from light in the pressing cassette, and covered with X-OMATBT film for 30 s to 3 min. The film was developed and imaged by the developer and fixing solution successively, dried and preserved, and then scanned. The strip density was quantified by densitometry using ImageJ software. The quantitative data were calculated by gray value, and β-actin was used as a control to analyze the relative protein expression. In addition, PI3K was used as the control for p-PI3K_Y607_, Akt was used as the control for p-Akt_Ser473_, and eNOS was used as the control for p-eNOS_Ser1177_.

### 2.10 Real-time fluorescence quantitative PCR (qRT-PCR)

Total RNA was extracted from the H9c2 cells, treated with TRIzol (Thermo Fisher Science), and premixed with the Takara (RR036A) reverse-transcription kit solvent. The RNA was incubated for 15 min at 37°C for reverse transcription and at 85°C for 5 s to inactivate the reverse transcriptase, and then cDNA was preserved at 4°C. Using TB Green PreMix Ex Tap II (Takara) and specific primers, real-time quantitative RT-PCR (Real-time Quantitative RT-PCR System, Thermo Fisher, USA) was used to quantify gene expression. The total reaction volume was 25 μL, comprising 10 μM for the final concentrations of specific primers, a 2 μL template, 2 × TB Green premix Ex Taq II, 50 × ROX reference fluorescent dye II, and enzyme-free sterile water. The amplification conditions were as follows: pre-denaturation at 95°C for 30 s,denaturation at 95°C for 5 s, annealing extension for 34 s at 60°C, for 40 cycles. We determined the threshold cycle (CT value) and relative gene expression by applying 2^−ΔΔCT^ and ΔΔCT=CT (β-actin) – CT (gene).

The primer sequences were as follows:

rat PI3K F, 5′-CTTGCCTCCATTCACCACCTCT-3′;

rat PI3K R, 5′-GCCTCTAATCTTCTCCCTCTCCTTCTC-3′;

rat Akt F, 5′-TGTCTCGTGAGCGCGTGTTTT-3′;

rat Akt R, 5′-CCGTTATCTTGATGTGCCCGTC-3′;

rat eNOS F, 5′-CCAGCTAGCCAAAGTCACCAT-3′;

eNOS R, 5′-GTCTCG GAGCCATACAGGATT-3′;

rat β-actin F, 5′-CACCACACCTTCTACAATGAGC-3′;

rat β-actin R, 5′-GTGATCTCCTTCTGCATCCTGT-3′.

All primers were synthesized by SangonBiotech (Shanghai, China).

### 2.11 Statistical analysis

We applied SPSS 26.0 software for data analysis. Measurement data that followed a normal distribution are expressed as x ± s. Single-factor analysis of variance was used for comparisons among multiple groups, with the LSD-t method used for post-hoc testing between groups. A significant difference was set at P < 0.05.

## 3. Results

### 3.1 Quality evaluation of SBP

To identify the main components of SBP, the retention time and ultraviolet spectrum (ESI, 2.5 kV, T350, 10 eV, 30 psi, 10 L/min, 100–1300 u) obtained by GC-MS were compared by high-performance liquid chromatography (203, 280 nm). Its ingredients included cinnamic acid, ginsenoside Re, cinnamic acid, cinnamaldehyde, ginsenoside Rb1, ginsenoside Rc, ginsenoside Rb2, ginsenoside Rb3, bufalin, ginsenoside Rd, cholic acid, ursodeoxycholic acid, hyodeoxycholic acid, cinnamic acid, chenodeoxycholic acid, and deoxycholic acid (Table 1; SBP components provided by Shanghai Hutchison Huangpu Pharmaceutical Co., Ltd., China) [29].

**Table 1.**
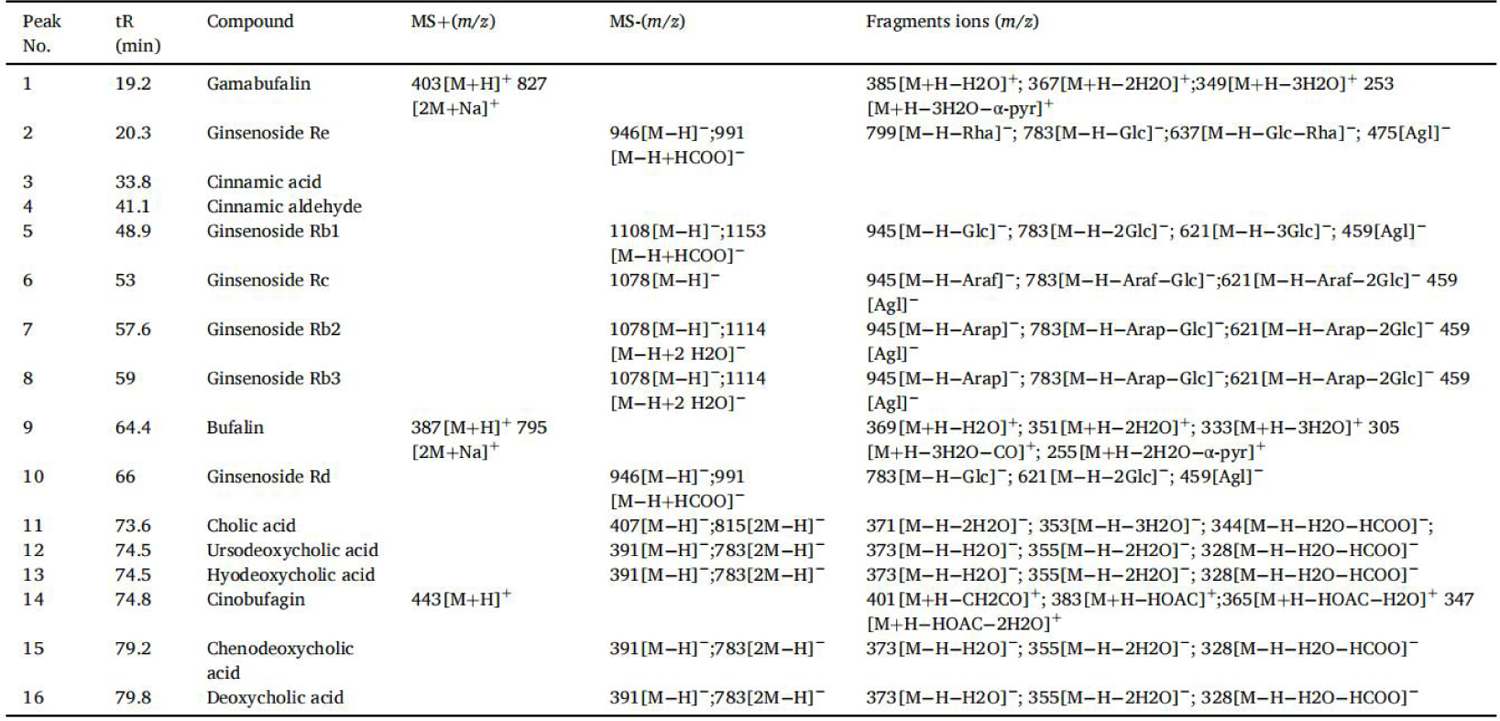
Mass spectral fragmentation results in HPLC/DAD.

### 3.2 Cell survival rate and apoptosis rate in each group

Compared with group 1, the cell survival rate decreased and the apoptosis rate increased in group 2 (P<0.05), indicating that H/R caused cardiomyocyte injury. Compared with group 2, the cell survival rate increased and the apoptosis rate decreased in group 3, indicating that SBP increased the cell survival rate and decreased the apoptosis rate and protected cardiomyocytes from H/R injury. Compared with group 3, cell viability decreased and the apoptosis rate increased in groups 5, 7, and 9, indicating that SBP increased the cell survival rate and decreased the apoptosis rate through the PI3K/Akt/eNOS signal pathway. When group 4 was compared with groups 5, 6, and 7, the cell survival rate increased and the apoptosis rate decreased (P<0.05). Compared with group 9, the apoptosis rate of group 8 decreased (P<0.05), and the cell survival rate increased but not significantly. When group 4 was compared with groups 5, 6, and 7 or 8 and 9, the addition of SBP increased the survival rate and decreased the apoptosis rate of PI3K/Akt/eNOS H/R cells, indicating that PI3K/Akt/eNOS inhibitors did not inhibit the protective effect of SBP on H/R cells. There may be other pathways by which SBP protects H/R cells.

**Table 2.** Comparison of cell survival rate and apoptotic rate among different groups.

| Group | cell survival rate (%)<br>(n=12) | Cell apoptosis rate (%)<br>(n=3) |
| --- | --- | --- |
| ①Control | $100 \pm 0$ | $4.04 \pm 1.60$ |
| ②H/R | $53.29 \pm 13.96^{acdf}$ | $34.16 \pm 1.55^{acdf}$ |
| ③H/R+SBP | $64.99 \pm 8.59^{begi}$ | $12.78 \pm 0.13^{begi}$ |
| ④H/R+LY294002 | $36.40 \pm 6.27^{be}$ | $42.55 \pm 0.63^{be}$ |
| ⑤H/R+LY294002+SBP | $53.85 \pm 6.18^{cd}$ | $29.99 \pm 0.53^{cd}$ |
| ⑥H/R+MK2206 | $35.38 \pm 10.64^{bg}$ | $38.57 \pm 1.54^{bg}$ |
| ⑦H/R+MK2206+SBP | $46.77 \pm 9.51^{cf}$ | $21.74 \pm 0.57^{cf}$ |
| ⑧H/R+L-NAME | $43.27 \pm 8.33$ | $35.16 \pm 1.99^i$ |
| ⑨H/R+L-NAME+SBP | $52.14 \pm 9.06^c$ | $16.35 \pm 0.38^{ch}$ |
| F | $38.157^{**}$ | $370.472^{**}$ |

Note:Note: H/R: hypoxia-reoxygenation model; SBP: Shexiang Baixin Pill; LY294002:PI3K inhibitor; MK2206:AKT inhibitor; L-NAME:eNOS inhibitor; \* $P<0.05$ , \*\* $P<0.01$ ; a and 1, b and 2、 c and 3、 d and 4、 e and 5、 f and 6、 g and 7、 h and 8、 I and 9 compared; $P<0.05$

**Figure 3.2.1.**
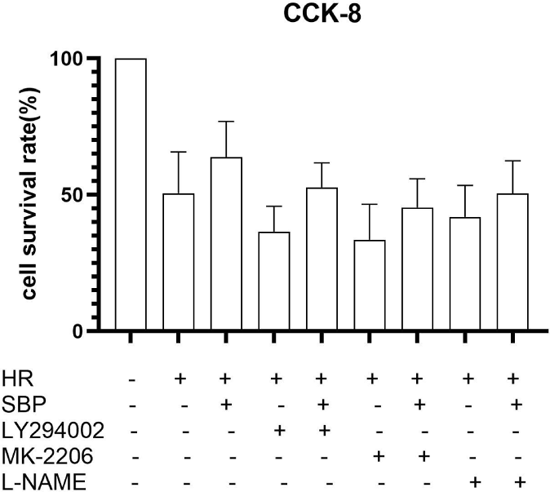
Comparison of cell survival rate among different groups.

**Figure 3.2.2.**
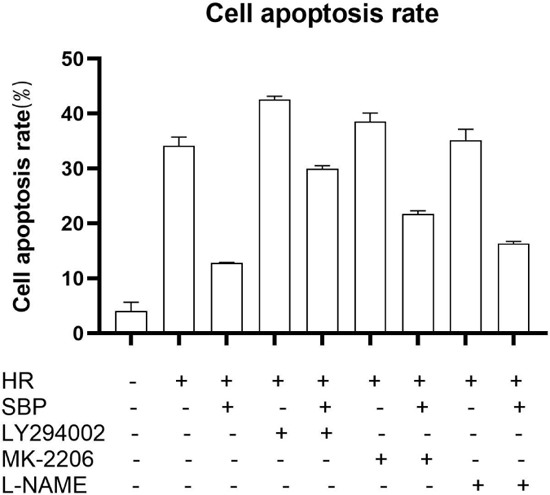
Comparison of apoptotic rate among different groups.

**Figure 3.2.3.**
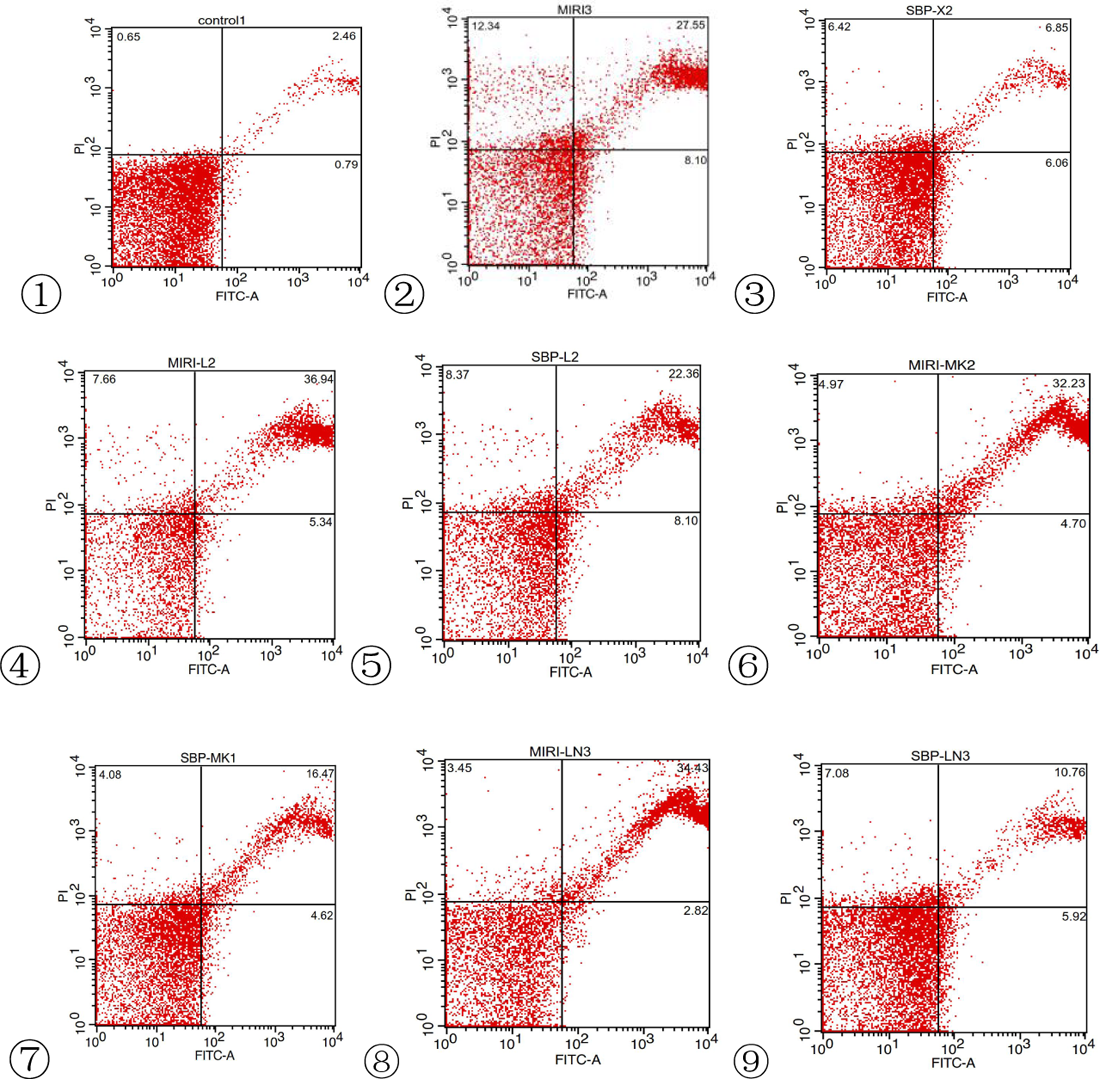
Flow cytometric detection map of cells in each group.

Conclusion: SBP increased the survival rate and decreased the apoptosis rate of hypoxia-reoxygenation (H/R) cardiomyocytes. The inhibition of the PI3K/Akt/eNOS signal pathway weakened the above-mentioned effects of SBP, indicating that the anti-apoptotic effect of SBP was achieved by activating the PI3K/Akt/eNOS-signaling pathway. The addition of PI3K/Akt/eNOS inhibitor did not completely inhibit the protective effect of SBP on H/R cardiomyocytes, suggesting that there may be other mechanisms by which SBP protected H/R cardiomyocytes.

### 3.3 p-PI3K/PI3K, p-Akt/Akt, and p-eNOS/eNOS protein phosphorylation levels in each group

The phosphorylation levels of p-PI3K_Y607_, p-Akt_Ser473,_ and p-eNOS_Ser1177_ in cardiomyocytes were detected by western blotting. The results showed that compared with group 2, the phosphorylation level of p-PI3K_Y607_, p-Akt_Ser473_, and p-eNOS_Ser1177_ increased in group 3 (P<0.05), indicating that SBP played a role in protecting cardiomyocytes from H/R injury. Compared with group 3, the phosphorylation levels of p-PI3K_Y607_, p-Akt_Ser473_, and p-eNOS_Ser1177_ in groups 5, 7, and 9 decreased significantly (P<0.05), indicating that SBP increased the phosphorylation of these proteins through the PI3K/Akt/eNOS signal pathway. There was no significant difference in protein phosphorylation levels of p-PI3K_Y607_, p-AktSer473, and p-eNOS_Ser1177_ between groups 4 and 5, 6 and 7, and 8 and 9, indicating that the inhibition of the PI3K/Akt/eNOS signal pathway did not decrease in spite of the addition of SBP,H/R, indicating that the effect of SBP on increasing the phosphorylation level of p-PI3K_Y607_, p-Akt_Ser473_, and p-eNOS_Ser1177_ protein occurred through activation of the PI3K/Akt/eNOS signal pathway.

**Table 3.**
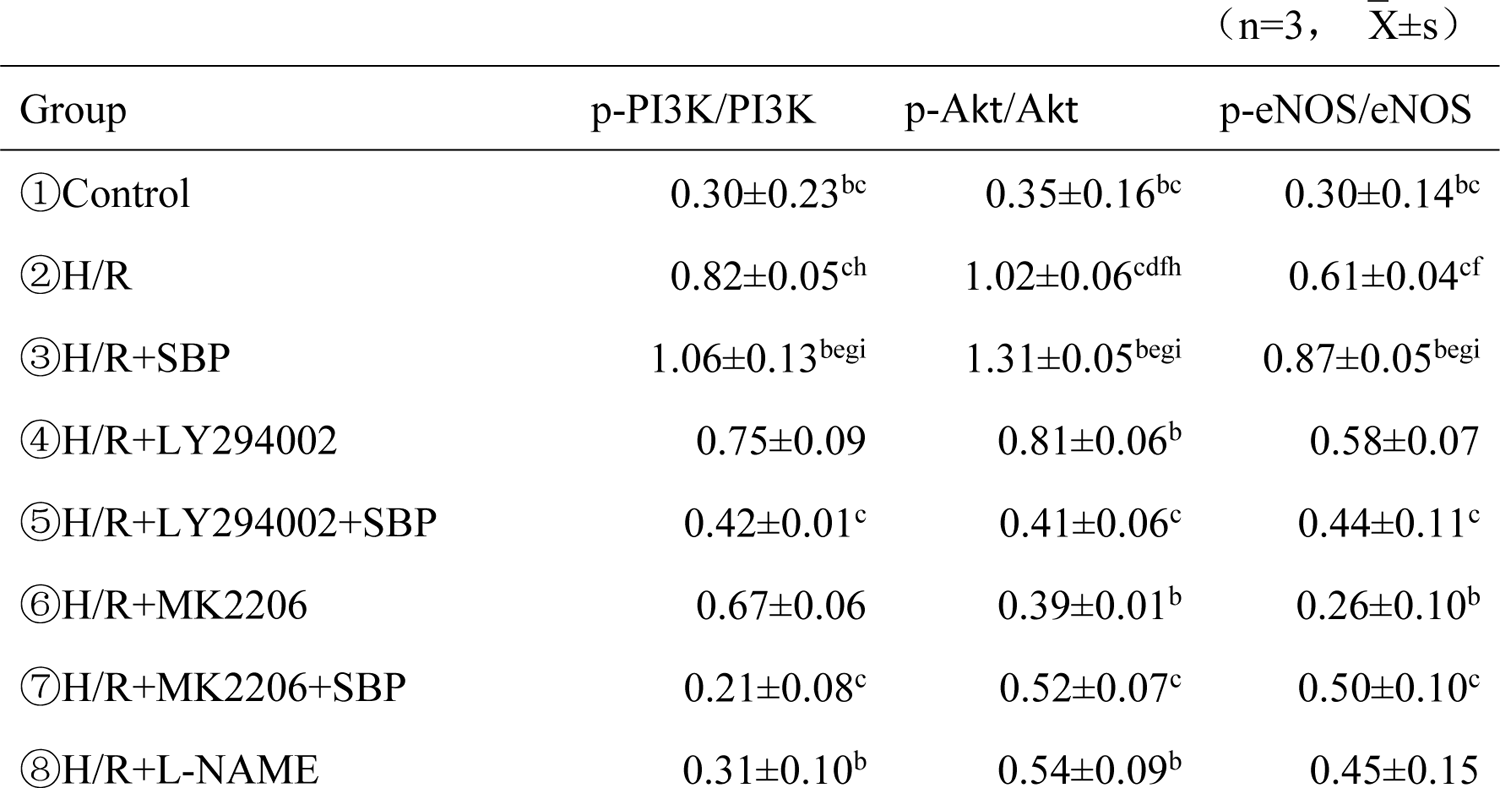
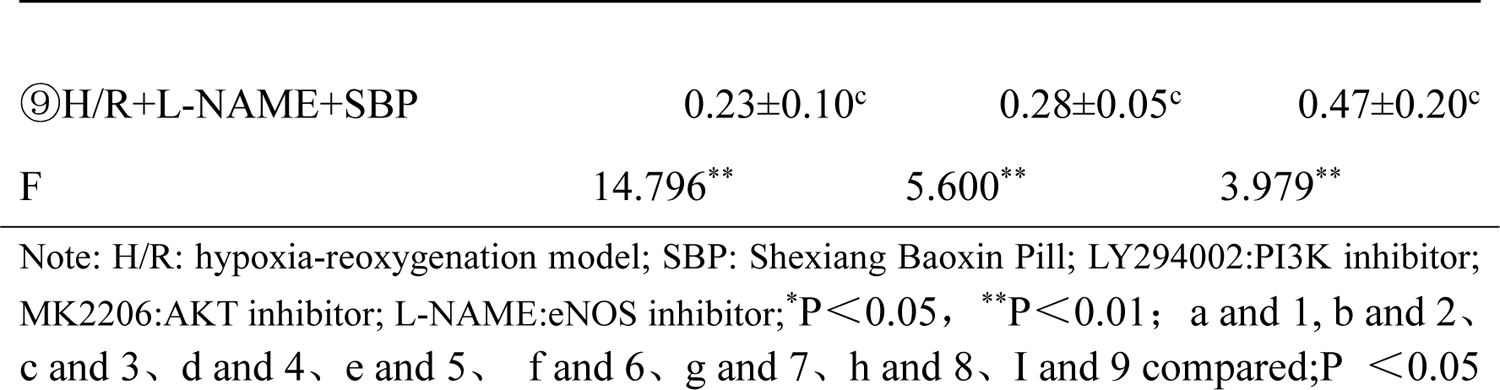
Comparison of p-PI3K/PI3K, p-Akt/Akt, and p-eNOS/eNOS protein expression in each group.

**Figure 3.3.1.**
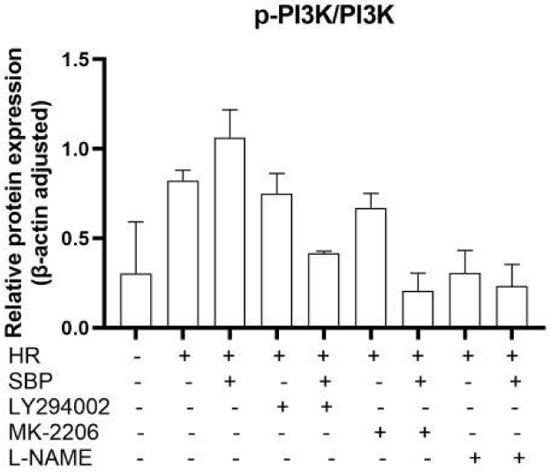
Proportion of p-PI3KY607 cell phosphorylation in each group.

**Figure 3.3.2.**
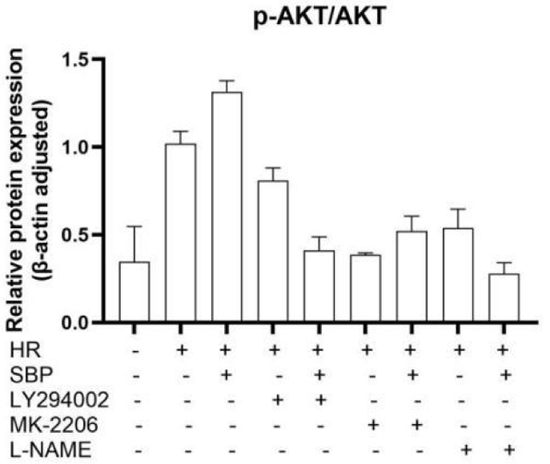
Proportion of p-AktSer473 cell phosphorylation in each group.

**Figure 3.3.3.**
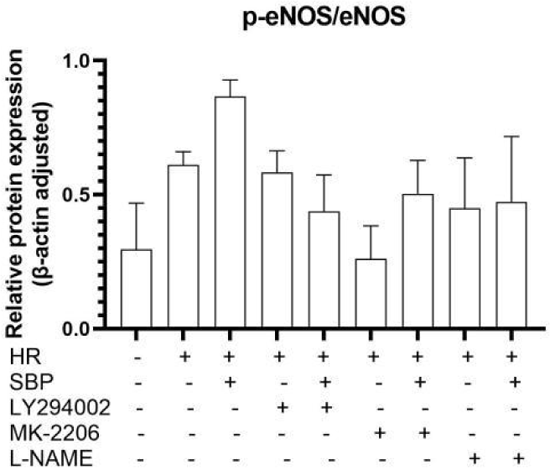
Proportion of p-eNOS_Ser1177_ cell phosphorylation in each group.

**Figure 3.3.4.**
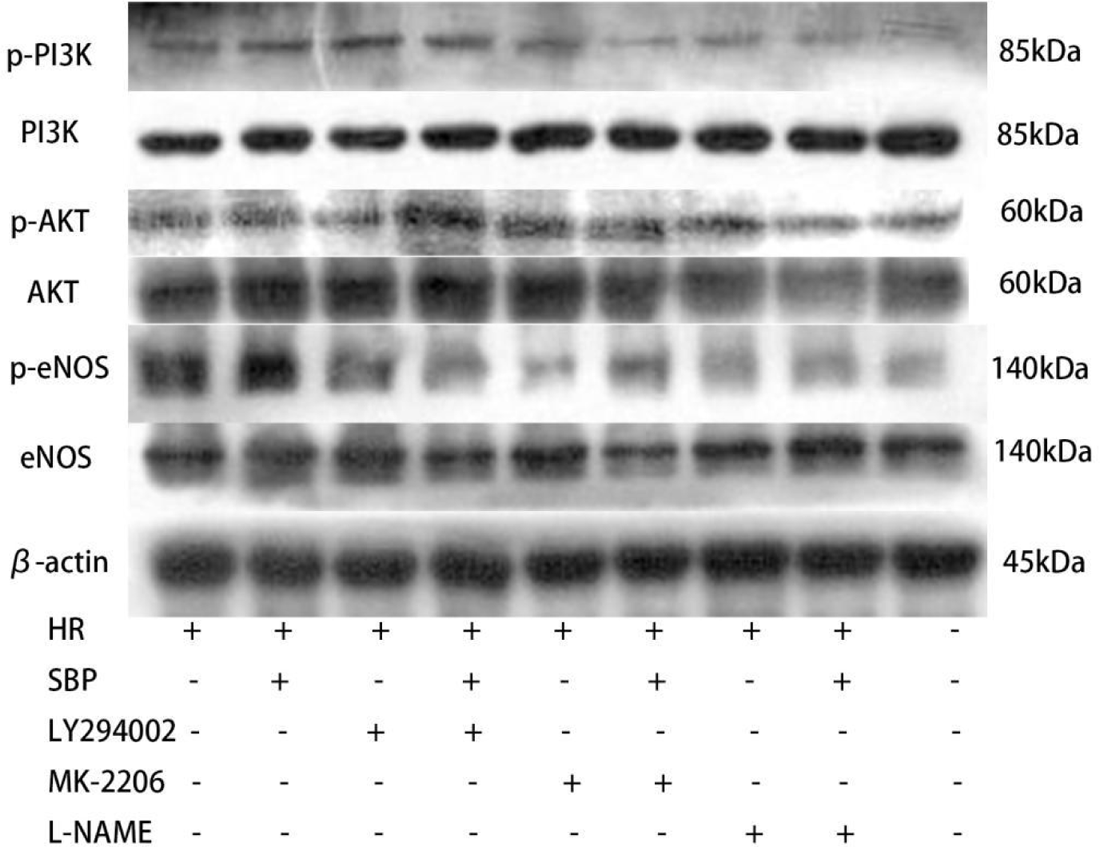
Comparison of relative expression levels of p-PI3KY607, p-AktSer473, and p-eNOSSer1177 cellular proteins in different groups.

Conclusion: SBP increased the level of p-PI3KY607, p-AktSer473, and p-eNOSSer1177 protein phosphorylation in H/R cardiomyocytes, and the inhibition of PI3K/Akt/eNOS signal pathway will attenuate the above-mentioned effects of SBP, indicating that SBP increases the phosphorylation of PI3K/Akt/eNOS signaling pathway proteins to protect against H/R injury.

### 3.4 PI3K, Akt, and eNOS mRNA levels in cells of each group

The relative expression of PI3K, Akt, and eNOS was observed by real-time fluorescence quantitative PCR. Compared with group 1, the relative expression of mRNA of PI3K, Akt, and eNOS increased in group 2, indicating that the PI3K/Akt/eNOS signal pathway was activated in H/R cardiomyocytes. Compared with group 2, the level of Akt mRNA in group 3 increased but not significantly, and the levels of PI3K and eNOS mRNA increased significantly (P<0.05), indicating that SBP increased the levels of PI3K, Akt, and eNOSmRNA in H/R cardiomyocytes. Compared with group 3, the relative expression of mRNA of PI3K, Akt, and eNOS in groups 5, 7, and 9 decreased significantly (P<0.05), indicating that the inhibition of PI3K/Akt/eNOS signal pathway significantly decreased the levels of PI3K and Akt eNOS mRNA in the three groups. Therefore, SBP played a protective role in H/R cardiomyocytes by activating the PI3K/Akt/eNOS pathway. There was no significant difference in the relative expression of mRNA of PI3K, Akt, and eNOS between groups 4 and 5, 6 and 7, and 8 and 9, indicating that after inhibiting the PI3K/Akt/eNOS signaling pathway, in spite of the addition of SBP, the mRNA levels of PI3K, Akt, and eNOS in H/R did not significantly increase. This indicated that the effect of SBP on increasing the mRNA levels of PI3K, Akt, and eNOS was achieved by activating the PI3K/Akt/eNOS signaling pathway.

**Table 4.**
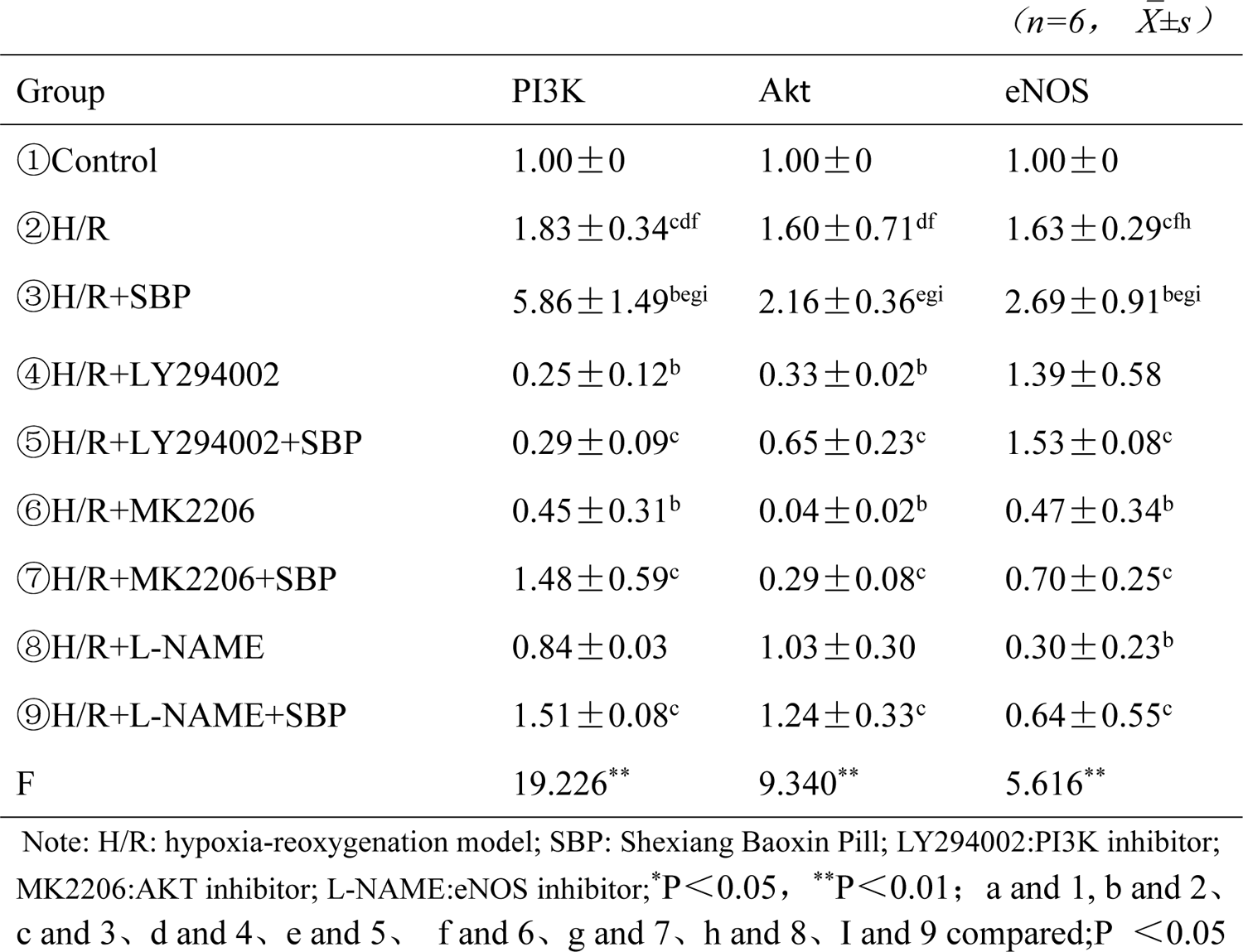
Comparison of PI3K, Akt, and eNOS mRNA levels of cells in each group.

**Figure 3.4.1.**
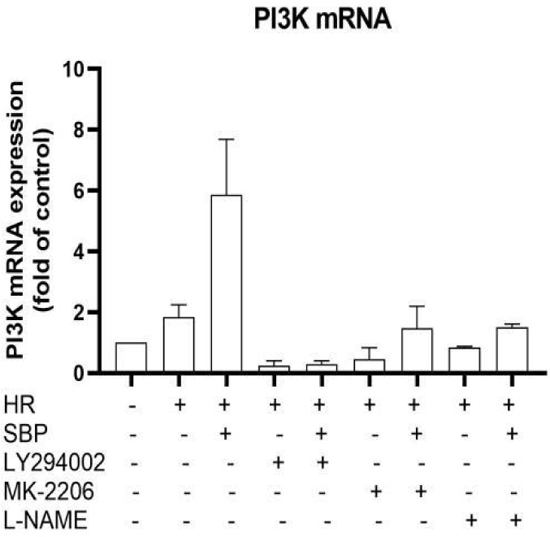
Comparison of PI3K mRNA levels of cells in different groups.

**Figure 3.4.2.**
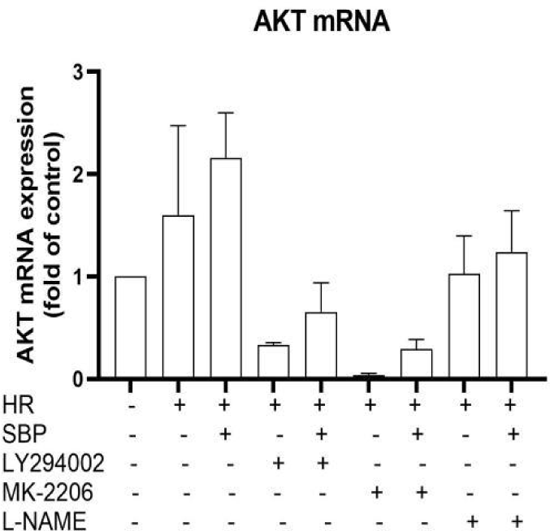
Comparison of Akt mRNA levels of cells in different groups.

**Figure 3.4.3.**
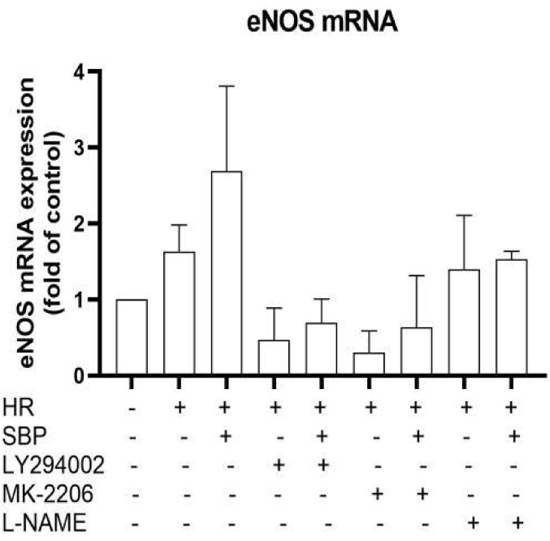
Comparison of eNOS mRNA levels of cells in different groups.

Conclusion: SBP increased the relative expression of mRNA of PI3K, Akt, and eNOS in H/R cardiomyocytes, and the inhibition of the PI3K/Akt/eNOS signal pathway attenuated the above-mentioned effects of SBP, indicating that SBP increased the relative expression of mRNA in the PI3K/Akt/eNOS signal pathway to protect against H/R injury.

### 3.5 Bax, Bcl-2, and Caspase-3 cell protein expression and Bcl-2/Bax ratio in each group

The protein expressions of Bax, Bcl-2, and caspase-3 in cardiomyocytes were detected by western-blot analysis. Compared with group 1, the expression of the apoptosis executive factor Caspase-3 increased and the ratio of Bcl-2/Bax decreased in group 2 (P<0.05), indicating that the expression of apoptosis-related proteins in cardiomyocytes increased and the ratio of Bcl-2/Bax decreased to promote apoptosis. Compared with group 2, the expression of anti-apoptotic factor Bcl-2 and the ratio of Bcl-2/Bax increased (P<0.05) in group 3, while the expression of pro-apoptotic factor Bax and apoptotic executive factor Caspase-3 decreased (P<0.05), indicating that SBP increased Bcl-2 and decreased the expression of Bax and Caspase-3, protecting cardiomyocytes and reducing apoptosis. Compared with group 3, there was no significant difference in the ratio of pro-apoptotic factors Bax and Bcl-2/Bax in groups 5, 7, and 9. There was no significant difference in the expression of anti-apoptotic factor Bcl-2 and apoptotic executive factor Caspase-3 among the five groups, but there was a significant difference in the expression of anti-apoptotic factor Bcl-2 and apoptotic executive factor Caspase-3 between groups 7 and 9 (P<0.05). The comparison between group 3 and 5, 7, and 9 showed that inhibition of the PI3K/Akt/eNOS signal pathway impacted the effect of SBP on increasing the ratio of Bcl-2 to Bcl-2/Bax and reducing Bax and Caspase-3. The inhibition of Akt/eNOS had a significant effect on Bcl-2 and Caspase-3, indicating that SBP played an important role in anti-apoptosis by activating the PI3K/Akt/eNOS signal pathway, and downstream Akt/eNOS had a significant effect on apoptosis. For group 4, there was no significant difference in the expression of anti-apoptosis factor Bcl-2, pro-apoptosis factor Bax, apoptosis executive factor Caspase-3, and the Bcl-2/Bax ratio between 4 and 5, 6 and 7, or 8 and 9, indicating that the inhibition of this pathway had no significant effect on the increase in the ratio of Bcl-2 and Bcl-2/Bax and the decrease in Bax and Caspase-3 protein expression even with the addition of SBP. It suggested that the anti-apoptotic effect of SBP was realized by activating the PI3K/Akt/eNOS signal pathway.

**Table 5.** Protein expression of Bax, Bcl-2, and caspase-3 in cells of each group.

| ($n=3, \bar{X} \pm s$) | | | | |
| --- | --- | --- | --- | --- |
| Group | Bax | Bcl-2 | Caspase-3 | Bcl-2/Bax |
| ①Control | $0.91 \pm 0.02$ | $0.73 \pm 0.08$ | $0.58 \pm 0.16$ | $0.81 \pm 0.09$ |
| ②H/R | $1.28 \pm 0.04^{cf}$ | $0.95 \pm 0.08^c$ | $1.33 \pm 0.07^{acd fh}$ | $0.75 \pm 0.07^{ac}$ |
| ③H/R+SBP | $1.02 \pm 0.05^b$ | $1.34 \pm 0.18^{bgi}$ | $1.02 \pm 0.01^{bgi}$ | $1.31 \pm 0.16^b$ |
| ④H/R+LY294002 | $0.98 \pm 0.10$ | $1.23 \pm 0.10$ | $0.74 \pm 0.13^b$ | $1.25 \pm 0.03$ |
| ⑤H/R+LY294002+SBP | $0.74 \pm 0.18$ | $1.11 \pm 0.14$ | $0.81 \pm 0.23$ | $1.63 \pm 0.57$ |
| ⑥H/R+MK2206 | $0.90 \pm 0.09^b$ | $1.13 \pm 0.11$ | $0.75 \pm 0.15^b$ | $1.28 \pm 0.23$ |
| ⑦H/R+MK2206+SBP | $1.08 \pm 0.12$ | $0.93 \pm 0.16^c$ | $0.65 \pm 0.10^c$ | $0.86 \pm 0.11$ |
| ⑧H/R+L-NAME | $0.90 \pm 0.09$ | $0.72 \pm 0.16$ | $0.64 \pm 0.05^b$ | $0.80 \pm 0.12$ |
| ⑨H/R+L-NAME+SBP | $0.80 \pm 0.15$ | $0.84 \pm 0.24^c$ | $0.56 \pm 0.08^c$ | $1.11 \pm 0.43$ |
| F | 3.883** | 4.275** | 7.728** | 2.580* |

Note: H/R: hypoxia-reoxygenation model; SBP: Shexiang Baixin Pill; LY294002:PI3K inhibitor; MK2206:AKT inhibitor; L-NAME:eNOS inhibitor; \* $P<0.05$ , \*\* $P<0.01$ ; a and 1, b and 2、c and 3、d and 4、e and 5、 f and 6、 g and 7、 h and 8、 I and 9 compared; $P <0.05$

**Figure 3.5.1.**
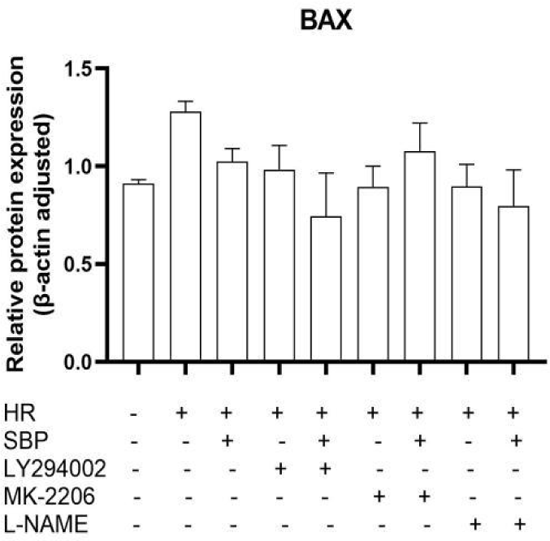
Comparison of Bax protein levels among different groups.

**Figure 3.5.2.**
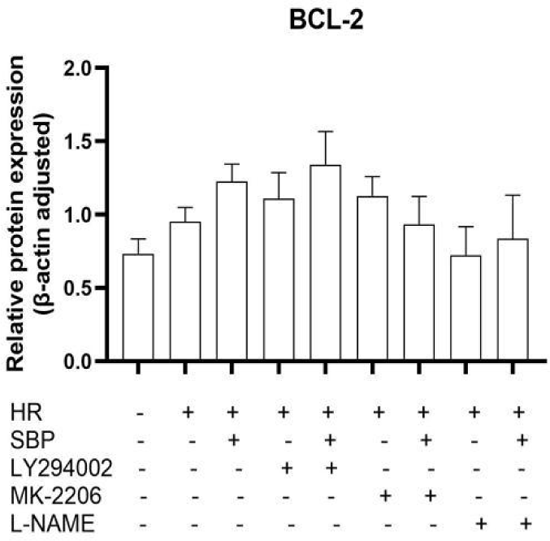
Comparison of Bcl-2 protein levels among different groups.

**Figure 3.5.3.**
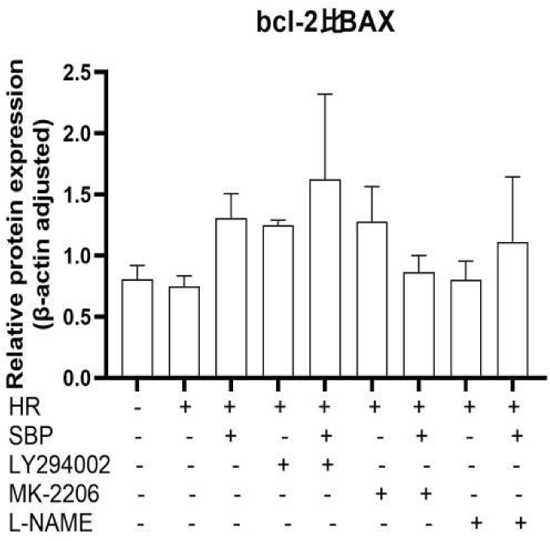
Comparison of Bcl-2/Bax protein levels among different groups.

**Figure 3.5.4.**
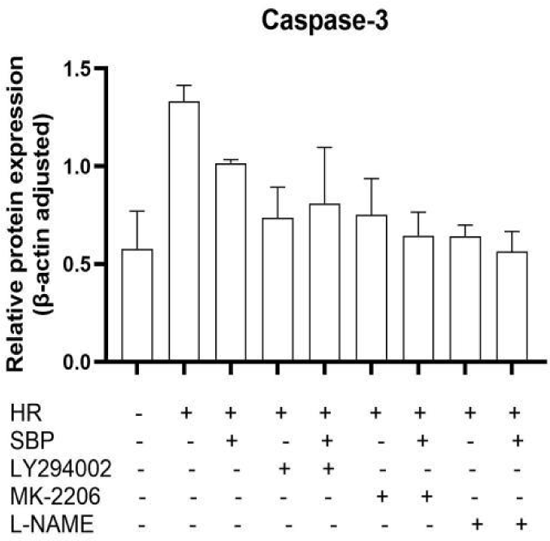
Comparison of caspase-3 protein levels among different groups.

**Figure 3.5.5.**
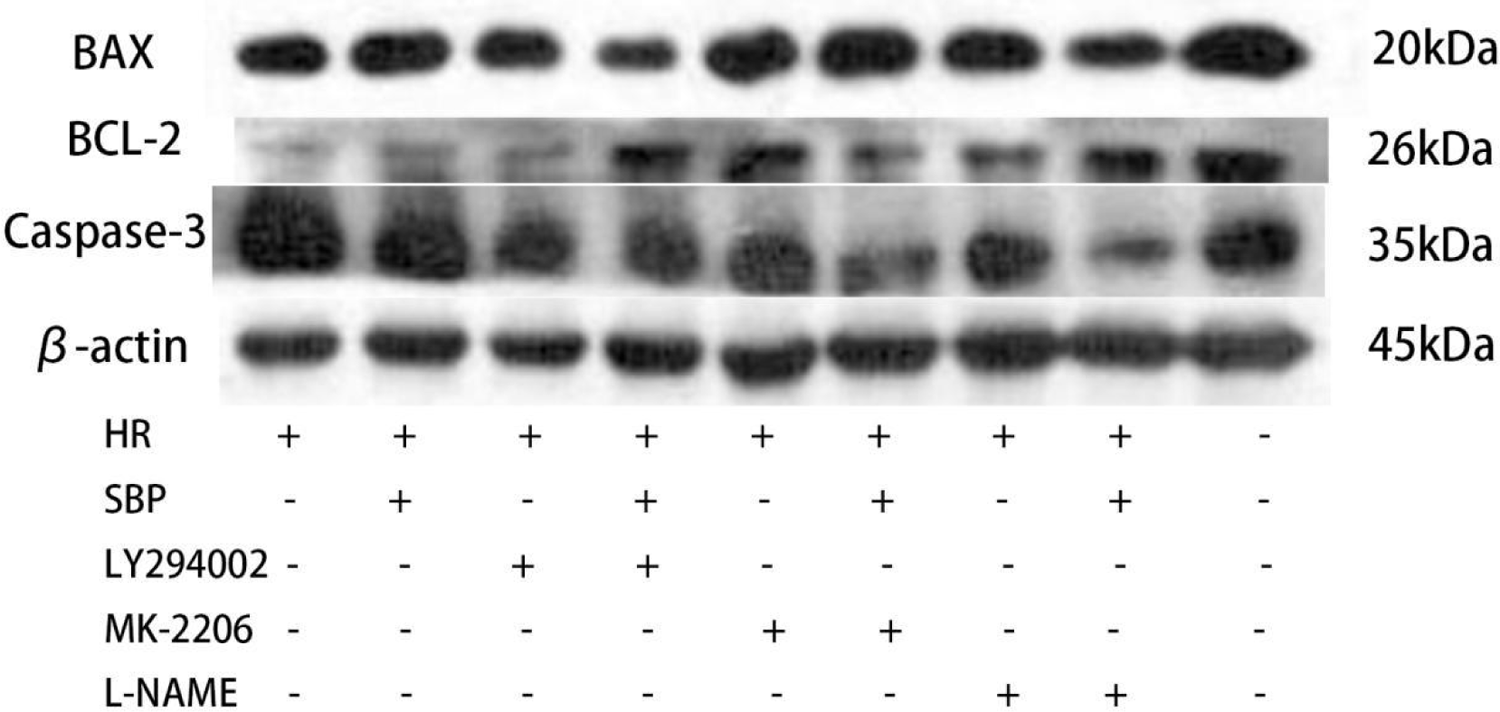
Comparison of Bax, Bcl-2, and caspase-3 protein levels among different groups.

Conclusion: SBP increased the ratio of Bcl-2 and Bcl-2/Bax in H/R cardiomyocytes and reduced Bax and Caspase-3 protein expression levels. Inhibition of the PI3K/Akt/eNOS signal pathway weakened the above effects of SBP, suggesting that SBP increased the ratio of Bcl-2 and Bcl-2/Bax in H/R cardiomyocytes through the PI3K/Akt/eNOS signal pathway, reduced the expression of Bax and Caspase-3 protein, and achieved anti-apoptotic protection against H/R injury.

### 3.6 Reactive oxygen species (ROS), gp91^phox^ protein, homocysteine (HCY), and malondialdehyde (MDA) cell expression in each group

The results showed that compared with group 1, the expression of ROS, gp91^phox^, HCY, and MDA increased in group 2, indicating that H/R increased the index of oxidative stress in cardiomyocytes. Compared with group 2, the expression of ROS, gp91^phox^, HCY, and MDA decreased in group 3 (P<0.05), indicating that SBP reduced the expression of ROS, gp91^phox^, HCY, and MDA and protected cardiomyocytes from oxidative stress. Compared with group 3, ROS increased and gp91^phox^ decreased significantly in group 5 (P<0.05), ROS increased and gp91^phox^ decreased but not significantly in groups 7 and 9, and the expression of HCY and MDA increased in groups 5, 7, and 9 (P<0.05), indicating that SBP significantly decreased the expression of HCY, MDA, ROS, and gp91^phox^ through the PI3K/Akt/eNOS signal pathway. When group 4 was compared with groups 5, 6, and 7 or 8 and 9, there was no significant difference in the expression of gp91^phox^ protein and HCY, but there was a significant difference in the expression of ROS and MDA (P<0.05). The results showed that after the inhibition of the PI3K/Akt/eNOS signal pathway, even if SBP was added, it had no significant effect on the expression of gp91^phox^ and HCY, but had a significant effect on the expression of ROS and MDA, indicating that the antioxidant stress of SBP was realized by activating this pathway. There may be other ways by which SBP reduces the expression of ROS and MDA.

**Table 6.** Expression of ROS, gp91^phox^ protein, HCY, and MDA in cells of each group.

( $\bar{X} \pm s$ )
| Group | ROS (n=15) | gp91<sup>phox</sup> (n=3) | HCY (pg/mL) (n=6) | MDA (nmol/mg) (n=6) |
| --- | --- | --- | --- | --- |
| ①Control | 65.93 ± 0.50 | 0.87 ± 0.22 | 7.78 ± 0.83 | 1.11 ± 0.18 |
| ②H/R | 121.41 ± 4.76<sup>acfh</sup> | 1.77 ± 0.18<sup>acfh</sup> | 20.03 ± 3.58<sup>ach</sup> | 2.09 ± 0.16<sup>acdf</sup> |
| ③H/R+SBP | 77.53 ± 1.43<sup>be</sup> | 1.22 ± 0.04<sup>b</sup> | 13.98 ± 2.33<sup>begi</sup> | 1.26 ± 0.22<sup>begi</sup> |
| ④H/R+LY294002 | 145.59 ± 7.86<sup>be</sup> | 1.41 ± 0.48 | 20.29 ± 1.62 | 2.43 ± 0.22<sup>be</sup> |
| ⑤H/R+LY294002+SBP | 90.56 ± 3.52<sup>cd</sup> | 0.95 ± 0.09 | 20.47 ± 1.40<sup>c</sup> | 1.84 ± 0.12<sup>cd</sup> |
| ⑥H/R+MK2206 | 110.15 ± 2.85<sup>bg</sup> | 0.85 ± 0.14<sup>b</sup> | 18.58 ± 1.49 | 2.59 ± 0.12<sup>bg</sup> |
| ⑦H/R+MK2206+SBP | 78.98 ± 1.11<sup>f</sup> | 0.77 ± 0.06 | 21.10 ± 1.36<sup>c</sup> | 2.04 ± 0.20<sup>cf</sup> |
| ⑧H/R+L-NAME | 110.24 ± 1.83<sup>bi</sup> | 1.01 ± 0.33<sup>b</sup> | 14.02 ± 4.61<sup>bi</sup> | 2.10 ± 0.29<sup>i</sup> |
| ⑨H/R+L-NAME+SBP | 85.17 ± 2.93<sup>h</sup> | 1.00 ± 0.32 | 17.10 ± 0.91<sup>ch</sup> | 1.69 ± 0.23<sup>ch</sup> |
| F | 96.053<sup>**</sup> | 3.316<sup>**</sup> | 17.280<sup>**</sup> | 36.514<sup>**</sup> |

Note: H/R: hypoxia-reoxygenation model; SBP: Shexiang Baoxin Pill; LY294002:PI3K inhibitor; MK2206:AKT inhibitor; L-NAME:eNOS inhibitor; \* $P < 0.05$ , \*\* $P < 0.01$ ; a and 1, b and 2、c and 3、d and 4、e and 5、 f and 6、 g and 7、 h and 8、 I and 9 compared; $P < 0.05$

**Figure 3.6.1.**
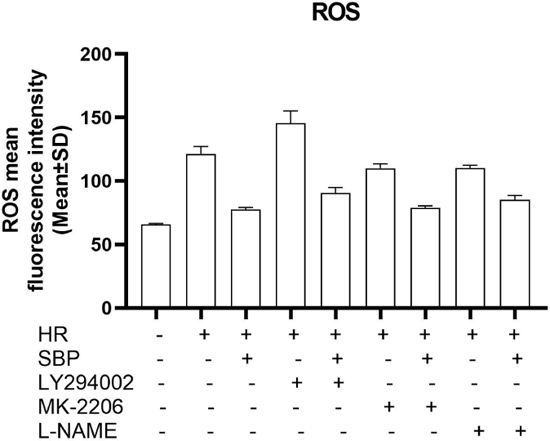
Comparison of ROS expression levels among different groups.

**Figure 3.6.2.**
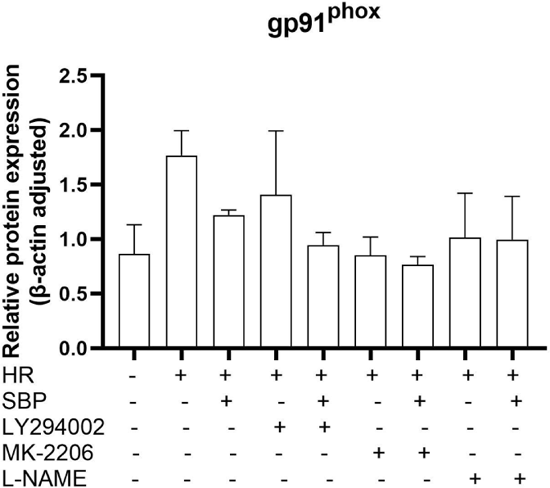
Comparison of gp91^phox^ protein expression levels among different groups.

**Figure 3.6.3.**
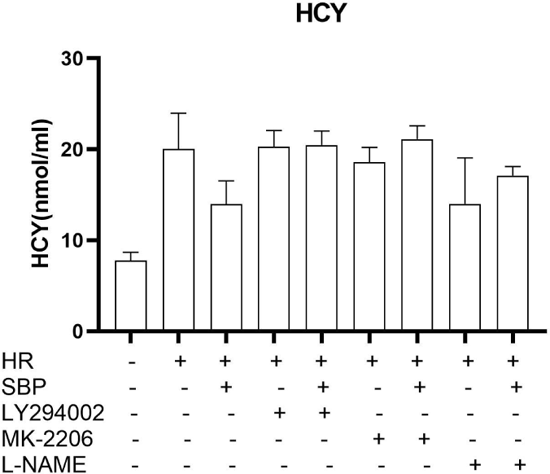
Comparison of HCY expression levels among different groups.

**Figure 3.6.4.**
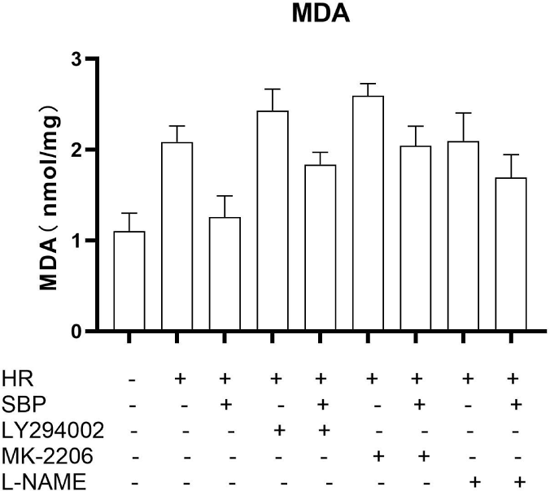
Comparison of MDA expression levels among different groups.

**Figure 3.6.5.**
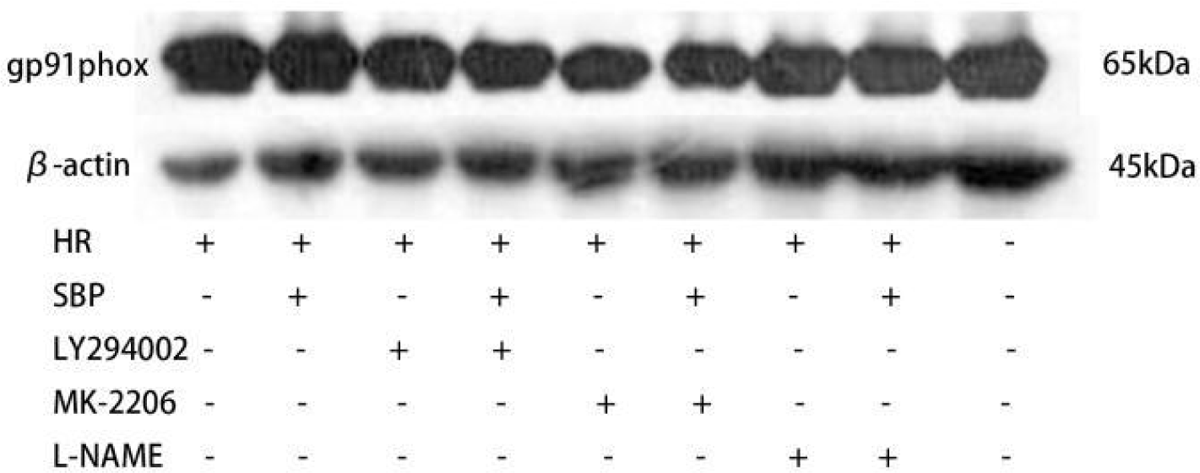
Comparison of gp91^phox^ protein expression levels among different groups.

**Figure 3.6.6.**
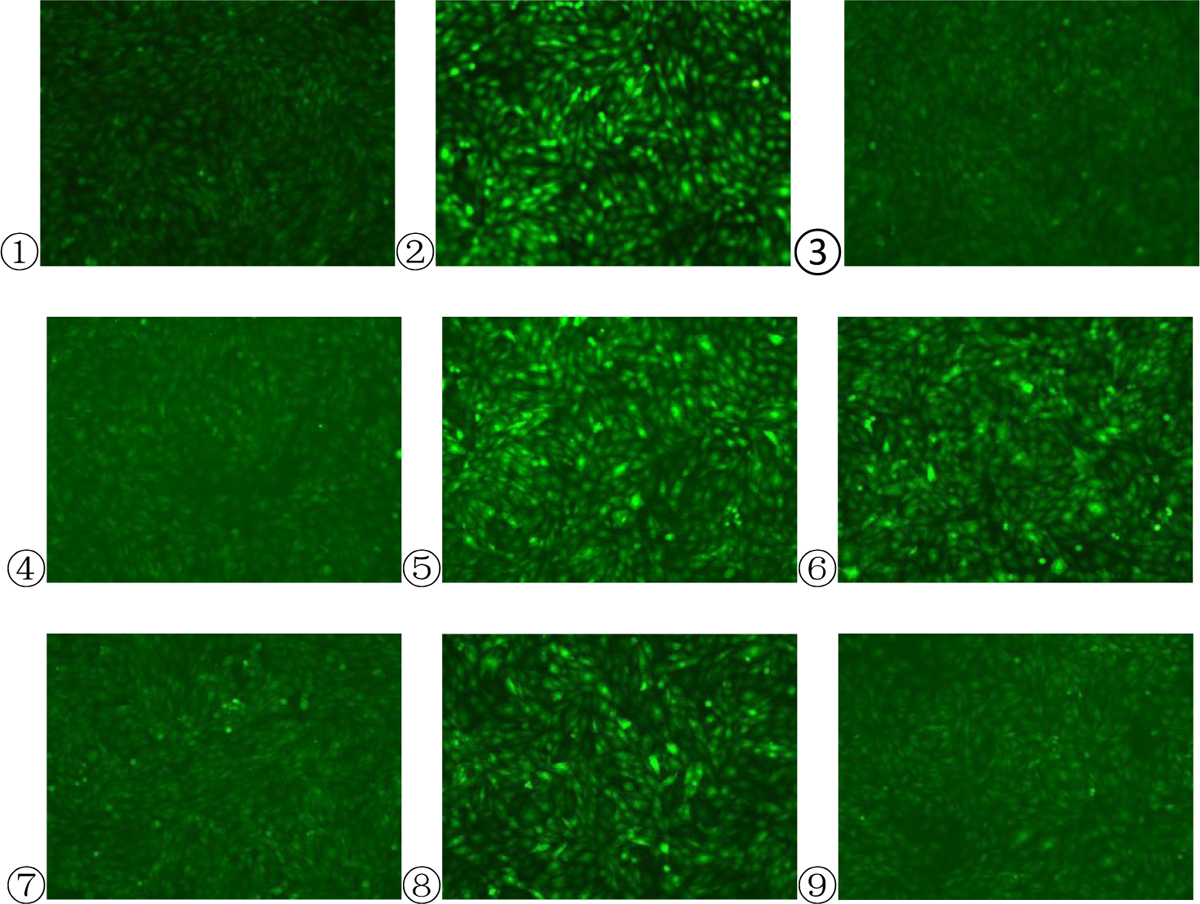
Comparison of ROS fluorescence staining expression among different groups.

Conclusion: SBP reduced the expression of ROS, gp91^phox^, HCY, and MDA in H/R cardiomyocytes, and the inhibition of the PI3K/Akt/eNOS signal pathway attenuated the above-mentioned effects of SBP, indicating that SBP reduced the expression of ROS, gp91^phox^, HCY, and MDA through the PI3K/Akt/eNOS signal pathway to achieve antioxidant stress protection against H/R injury and reduced the expression of ROS and MDA via other mechanisms.

### 3.7 Intracellular tumor necrosis factor-α (TNF-α), interleukin-6 (IL-6), and interleukin-18 (IL-18) content of each group

The contents of TNF-α, IL-6, and IL-18 in group 2 were higher than those in group 1 (P<0.05), indicating that H/R increased the content of inflammatory factors in cardiomyocytes. Compared with group 2, the contents of TNF-α, IL-6, and IL-18 decreased in group 3 (P<0.05), indicating that SBP decreased the expression of TNF-α, IL-6, and IL-18 and protected cardiomyocytes from the inflammatory response to H/R. Compared with group 3, the contents of TNF-α, IL-6, and IL-18 in groups 5, 7, and 9 increased, and the contents of IL-6 and IL-18 in groups 5 and 7 increased significantly. The results indicated that inhibition of this pathway affected SBP’s reduction of TNF-α, IL-6, and IL-18 levels and had a significant effect on IL-6 and IL-18, indicating that SBP played an anti-inflammatory role in H/R by activating this pathway. When comparing group 4 with groups 5, 6, and 7 or 8 and 9, the contents of TNF-α, IL-6, and IL-18 decreased but there was no significant difference, indicating that after inhibition of the PI3K/Akt/eNOS signal pathway, the addition of SBP had no significant effect on the reductions of TNF-α, IL-6, and IL-18 levels in H/R, suggesting that the anti-inflammatory effect of SBP was achieved by activating this pathway.

**Table 7.** Comparison of intracellular TNF-α, IL-6, and IL-18 contents in different groups.

( $n=4$ , $\bar{X} \pm s$ )
| Group | TNF-α (pg/mL) | IL-6 (pg/mL) | IL-18 (pg/mL) |
| --- | --- | --- | --- |
| ①Control | $3.04 \pm 1.59$ | $0.16 \pm 0.03$ | $104.43 \pm 4.19$ |
| ②H/R | $7.28 \pm 3.47^{ac}$ | $0.25 \pm 0.02^{ac}$ | $155.54 \pm 20.49^{ac}$ |
| ③H/R+SBP | $3.52 \pm 1.87^b$ | $0.15 \pm 0.04^{beg}$ | $109.27 \pm 9.23^{beg}$ |
| ④H/R+LY294002 | $5.45 \pm 0.54$ | $0.26 \pm 0.03$ | $140.09 \pm 24.23$ |
| ⑤H/R+LY294002+SBP | $3.58 \pm 0.72$ | $0.21 \pm 0.01^c$ | $168.58 \pm 14.47^c$ |
| ⑥H/R+MK2206 | $6.01 \pm 2.63$ | $0.22 \pm 0.05$ | $147.43 \pm 27.09$ |
| ⑦H/R+MK2206+SBP | $4.16 \pm 0.24$ | $0.22 \pm 0.02^c$ | $180.05 \pm 15.13^c$ |
| ⑧H/R+L-NAME | $7.12 \pm 1.32$ | $0.29 \pm 0.07$ | $164.55 \pm 26.95$ |
| ⑨H/R+L-NAME+SBP | $5.89 \pm 0.81$ | $0.26 \pm 0.04$ | $133.62 \pm 27.29$ |
| F | 2.428* | 4.369** | 4.820** |

Note: H/R: hypoxia-reoxygenation model; SBP: Shexiang Baoxin Pill; LY294002:PI3K inhibitor; MK2206:AKT inhibitor; L-NAME:eNOS inhibitor; \* $P < 0.05$ , \*\* $P < 0.01$ ; a and 1, b and 2、c and 3、d and 4、e and 5、 f and 6、 g and 7、 h and 8、 I and 9 compared; $P < 0.05$

**Figure 3.7.1.**
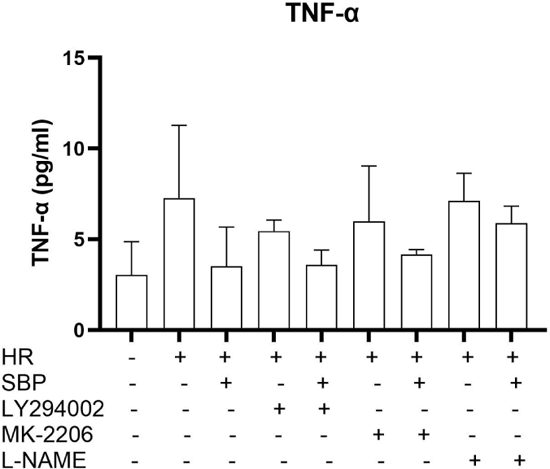
Comparison of TNF-α expression among different groups.

**Figure 3.7.2.**
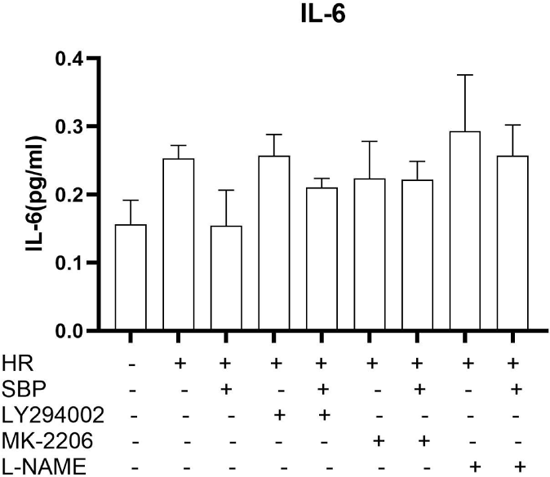
Comparison of IL-6 expression in each group.

**Figure 3.7.3.**
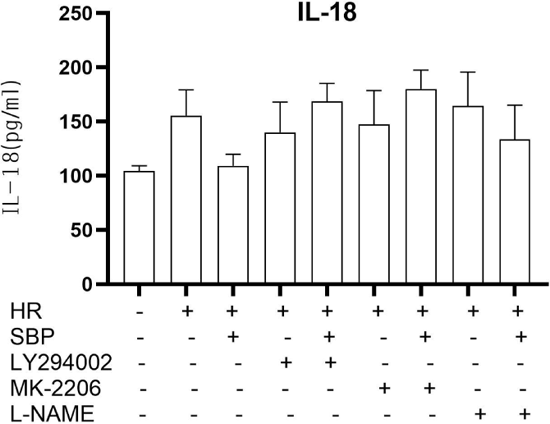
Comparison of IL-18 expression in each group.

Conclusion: SBP reduced the contents of TNF-α, IL-6, and IL-18 in H/R cardiomyocytes, and the inhibition of PI3K/Akt/eNOS signal pathway weakened the above effects of SBP, indicating that SBP reduced the contents of TNF-α, IL-6, and IL-18 through the PI3K/Akt/eNOS signal pathway.

## 4. Discussion

SBP exerted effects on antioxidant stress, anti-apoptosis, and the anti-inflammatory response by activating the PI3K/Akt/eNOS-signaling pathway and attenuating cardiomyocyte injury induced by H/R, resulting in an improved cell survival rate of H/R cardiomyocytes. Inhibition of PI3K, Akt, and eNOS impeded the increase in the cell survival rate induced by SBP, and SBP increased the level of p-PI3K_Y607_, p-Akt_Ser473_, and p-eNOS_Ser1177_ protein phosphorylation and mRNA in H/R cardiomyocytes. Inhibition of PI3K, Akt, and eNOS impeded the increase in p-PI3K_Y607_, p-Akt_Ser473_, p-eNOS_Ser1177_ protein phosphorylation, and mRNA levels of cardiomyocytes induced by SBP, while SBP increased the level of Bcl-2 protein and Bcl-2/Bax ratio and decreased the rate of apoptosis and the expression of Bax and Caspase-3. Inhibition of PI3K, Akt, and eNOS impeded the increase in Bcl-2 protein and Bcl-2/Bax ratio, the decrease in Bax and Caspase-3 expression, and the decrease in oxidative stress indices (ROS, HCY, MDA, and gp91^phox^) and inflammatory factors (TNF-α, IL-6, IL-18). Inhibition of PI3K, Akt, and eNOS inhibited the decrease in oxidative stress and inflammatory response.

MIRI is a common pathophysiological injury in STEMI patients [30], which can increase the area of myocardial infarction and adversely affect patient prognosis [4]. A compound preparation of traditional Chinese medicine, SBP can reduce MIRI and myocardial infarction area[21]. Previous studies have suggested that the PI3K/Akt/eNOS signal pathway is an important survival pathway in the MIRI signal transduction pathway, which plays an important role in inhibiting apoptosis and promoting cell survival [31].

Apoptosis has been regarded as a strictly controlled and perfect regulatory process, while cell necrosis has been regarded as an uncontrolled mode of cell death [32]. Budhram-Mahadeo et al. found that Caspase-3 and Bax are rapidly activated in MIRI after coronary artery occlusion; however, the anti-apoptotic protein Bcl-2 remains inactive or significantly decreased [33–34]. We found that SBP increased the survival rate of H/R cardiomyocytes and reduced the apoptosis rate, reduced the expression of Bax and Caspase-3 protein, increased the expression of Bcl-2 protein, and achieved an anti-apoptosis effect by activating the PI3K/Akt/eNOS signal pathway. Our results are consistent with their results. We also confirmed that the anti-apoptotic effect of SBP on H/R cardiomyocytes is related to the increase in the Bcl-2/Bax ratio. Chi et al. found that ibuprofen up-regulates the Bcl-2/Bax ratio through an Akt-related signaling pathway to reduce the occurrence of apoptosis in MIRI [35]. Therefore, increasing the Bcl-2/Bax ratio is the key target of anti-apoptosis in MIRI [36–37]. Syed Abd Halim et al. noted that apoptosis occurs through internal or external stimulation, which mainly involves mitochondrial damage and caspase-9 activation. Caspase-9 activates caspases-3 and −7, while the external pathway is mediated by the death receptor Bcl-2/Bax, resulting in an apoptosis cascade reaction. In short, the regulation of cardiomyocyte apoptosis is the key outcome of MIRI, which can directly lead to whether cardiac function will be permanently damaged. We confirmed that SBP reduced MIRI by activating the PI3K/Akt/eNOS signal pathway and anti-apoptotic effect.

Oxidative stress occurs in the early stage of reperfusion therapy in MIRI. A large number of oxidative stress factors such as ROS, gp91^phox^, HCY, and MDA are highly expressed, which leads to cell death. This study confirmed that SBP reduced the expression of these oxidative stress factors in H/R cardiomyocytes. Li et al. found that ginsenosides, the main component of SBP, reduce the oxidative damage factor ROS[40] and downregulate the expression of gp91^phox^ protein to reduce MIRI in a MIRI rat myocardial infarction model [41]. In our experiment, the aqueous extract of SBP was used, and some of the research results were consistent with the above results, indicating that SBP has high efficiency and multi-efficacy related to retaining the function of monomer components. At present, the increase in the serum Hcy level is an independent risk factor for cardiovascular disease [42], but there are few studies on the relationship between Hcy and MIRI. The main cause of Hcy-induced toxicity is oxidative stress, which primarily includes the oxidative production of Hcy to O_2_^-^ damage antioxidant system, and the covalent modification of proteins to homocysteine thiolactone after methylthio-tRNA synthetase is transformed into homocysteine thiolactone by a misediting mechanism. This leads to the formation of toxic protein oligomers, which directly damage the function of mitochondria and many other membranous organelles [42]. We therefore measured the total amount of Hcy to evaluate whether it caused MIRI and aggravated oxidative stress. SBP attenuated ROS-induced cardiac myocyte injury in a rat MIRI model by reducing Hcy. Our experiments confirmed that SBP reduced the degree of oxidative stress in MIRI by inhibiting Hcy through the PI3K/Akt/eNOS signal pathway, which expanded the novel idea that SBP exerted a multi-target effect. In addition, disulfides formed by Hcy self-oxidation not only induce ROS production [44–45] and increase the expression of NADPH oxidase 4, but Hcy has also been shown to change cell signal transduction by affecting the Akt/eNOS signal transduction pathway, aggravating oxidative stress, and reducing cardiomyocyte injury [46]. Hcy also decreases the ratio of Bcl-2/Bax, affects the mitochondrial membrane potential, leads to mitochondrial apoptosis, activates caspase-3, and causes irreversible damage to cardiomyocytes[45,47–48]. Therefore, it is very important to explore the role of SBP and Hcy in MIRI. In addition, Hcy is related to the promotion of lipid peroxidation by ROS. MDA is a common marker of lipid peroxidation induced by ROS. Low levels of MDA and ROS have shown a strong antioxidant capacity. This study confirmed that SBP decreased the expression of MDA and ROS in MIRI cardiomyocytes, suggesting that SBP may have an effect on anti-oxidative stress.

The inhibition of the inflammatory response can reduce myocardial structural and functional damage caused by the inflow of inflammatory factors into ischemic myocardial tissue [49–50]. SBP inhibits the release of inflammatory factors, which is a key target to reduce MIRI. We have confirmed that SBP can inhibit the inflammatory response through the PI3K/Akt/eNOS signal pathway, reduce the release of inflammatory factors TNF-α, IL-6, and IL-18, and achieve myocardial protection against H/R injury. We also confirmed that SBP can reduce the release of IL-18 from H/R cardiomyocytes through the PI3K/Akt/eNOS signal pathway. IL-18 is a key cytokine in pyroptosis [51]. It further magnifies the inflammatory cascade by inducing additional cytokines, adhesion molecules, and chemokines. As a result, the damaged myocardium releases various damage-related molecules such as ROS, thus activating inflammatory bodies, promoting the maturation of TNF-α and IL-18 precursors, and aggravating the occurrence of the inflammatory response [52]. Among them, pyroptosis is a pro-inflammatory form of programmed cell death, which is characterized by the formation of pores in the plasma membrane leading to cell swelling, cell membrane rupture [53], and promoting the release of TNF-α, IL-6, and IL-18 [54]. Therefore, SBP has the effect of anti-pyrolysis of MIRI cardiomyocytes. Saemann et al. confirmed that the increase in TNF-α and IL-6 can induce the formation of ROS in endothelial cell mitochondria [55], and ROS can stimulate the production of IL-18 in MIRI [56–57], promote apoptosis and calcium overload, lead to cardiomyocyte injury [58–59], and eventually lead to apoptosis and pyroptosis [60]. It has been confirmed that SBP can inhibit the production of ROS and the release of TNF-α, IL-6, and IL-18 in MIRI, indicating that SBP has a synergistic effect on inhibiting oxidative stress as well as the inflammatory response in MIRI. In addition, SBP activates the PI3K/Akt/eNOS signal pathway to increase the release of nitric oxide (NO), while NO has been shown to inhibit the release of IL-18 from macrophages exhibiting an anti-inflammatory effect [61]. These results suggest that SBP may reduce the inflammatory reaction and pyroptosis of MIRI.

This study possessed some limitations. We confirmed that SBP targets multiple sites in the treatment of H/R cardiomyocyte injury, but only at the cellular level and not in animal experiments. We have not yet demonstrated a role for specific components of SBP and the interaction between them and have not yet fully explored all the underlying mechanisms and signaling pathways involved in the treatment of MIRI with SBP. In terms of apoptosis, our experiment only showed the important factors in the pathway, and more comprehensive follow-up studies are therefore needed. In terms of oxidative stress, although many oxidative stress factors were analyzed in this experiment, the interaction of these factors and specific signal-transduction pathways needs to be clarified. With respect to inflammatory responses, although our experimental results revealed that the inhibition of inflammatory responses and pyroptosis were potential targets for the treatment of MIRI, this still needs to be confirmed by evaluating a variety of other signaling pathways and targets.

**Figure 4.**
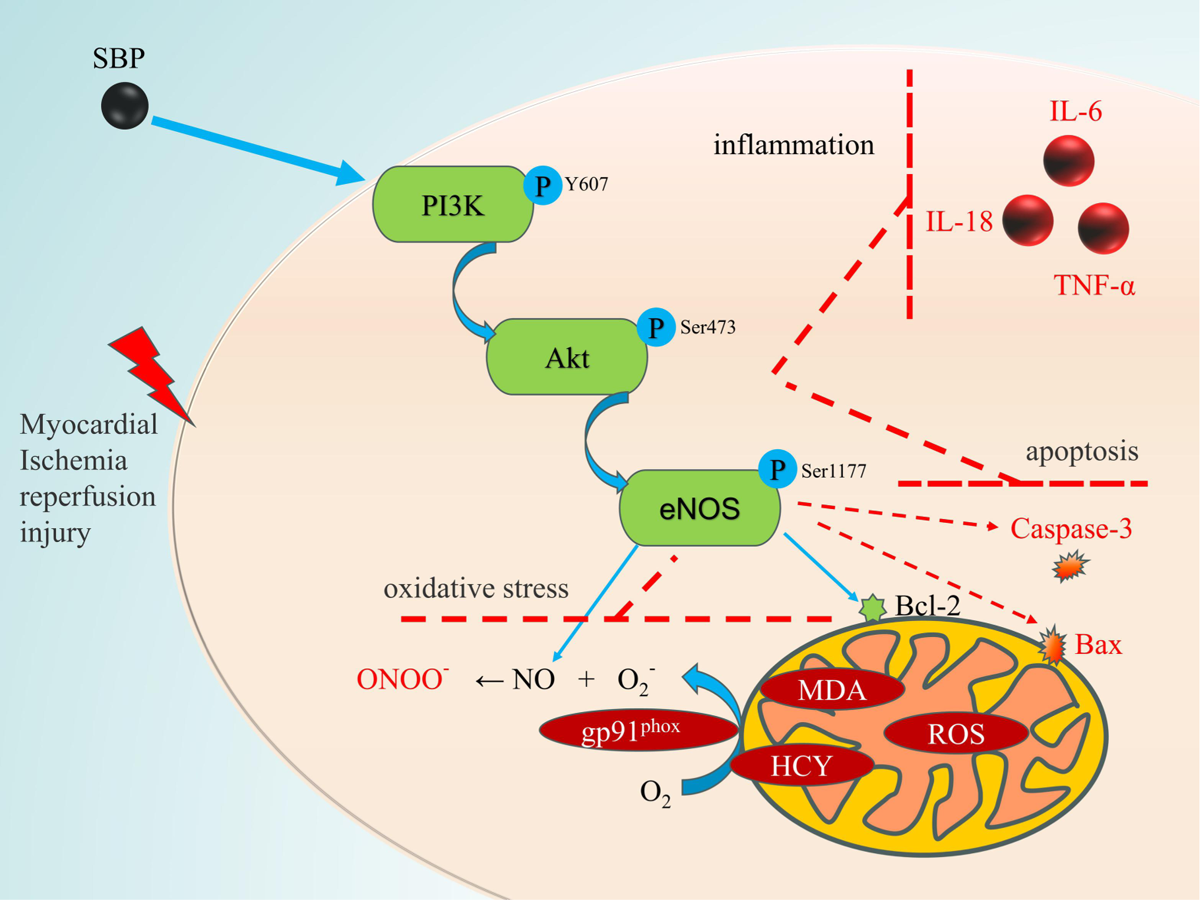
Mechanism underlying MIRI reduction in cardiomyocytes by SBP.

## 5. Conclusions

In summary, we ascertained that SBP exerts a protective effect on H9c2 cardiomyocytes exposed to hypoxia and reoxygenation through the PI3K/Akt/eNOS-signaling pathway and that it may inhibit apoptosis, oxidative stress, and inflammatory responses. SBP may additionally inhibit hypoxia and reoxygenation injury of cardiomyocytes and protect cardiomyocytes through multi-target effects. These findings are critical to further understanding SBP in the treatment and prevention of cardiovascular disease.

